# Neither fifty percent slow-wave sleep suppression nor fifty percent rapid eye movement sleep suppression does impair memory consolidation

**DOI:** 10.1101/2024.08.19.607534

**Authors:** Yulia V. Ukraintseva, Konstantin A. Saltykov, Olga N. Tkachenko

## Abstract

Establishing well-defined relationships between sleep features and memory consolidation is essential in comprehending the pathophysiology of cognitive decline commonly seen in patients with insomnia, depression, and other sleep-disrupting conditions.

Twenty-eight volunteers participated in two experimental sessions: a session with selective SWS suppression during one night and a session with undisturbed night sleep (as a control condition). Fifteen of them also participated in a third session with REM suppression. Suppression was achieved by presenting an acoustic tone. In the evening and the morning, the participants completed procedural and declarative memory tasks and the Psychomotor vigilance task (PVT). Heart rate variability analysis and salivary cortisol were used to control possible stress reactions on sleep interference.

SWS and REM suppression led to more than 50 percent reduction in amount of these stages. Neither vigilance nor memory consolidation was impaired after SWS or REM suppression. Unexpectedly, a beneficial effect of selective SWS suppression on PVT performance was found. Similarly, after a night with SWS suppression, the overnight improvement in procedural skills was higher than after a night with REM suppression and after a night with undisturbed sleep.

Our data brings into question the extent to which SWS and REM are truly necessary for effective memory consolidation to proceed. Moreover, SWS suppression may even improve the performance of some tasks, possibly by reducing sleep inertia associated with undisturbed sleep.

**Highlights:**

Our data brings into question the extent to which SWS and REM are truly necessary for effective memory consolidation to proceed.
Provided that sleep disturbances do not cause stress, half the usual amount of SWS or REM is sufficient for procedural and declarative memory consolidation.
Moreover, SWS suppression may even improve the performance of psychomotor vigilance task and finger sequence tapping task, possibly by reducing sleep inertia associated with undisturbed night sleep.

## 1. Introduction

The effect of sleep on human memory has gained significant attention in neuroscience research over the last three decades, resulting in a large body of studies (for review, see Rasch, Born, 2013; Brodt et al.,2023). Establishing well-defined relationships between sleep features and memory consolidation is of high clinical significance, contributing to a better understanding of the pathophysiology of memory impairment commonly seen in insomnia, depression, and other conditions characterized by disrupted sleep. However, a consensus among the researchers has not been achieved yet, as evidenced by the variety of conflicting theories and concepts (Rasch, Born, 2013; Poe, 2017; Sara, 2017).

How sleep following learning can enhance subsequent memory is still not clear. Even the actual impact of sleep on memory strength is still under considerable debate because the correlations observed between the sleep duration and architecture (the amount of SWS and REM, number of sleep cycles, etc.) and memory consolidation are often not replicated across studies (Casey et al., 2016; Genzel et al., 2009; Lahl et al., 2008; Backhaus, Junghanns, 2006; Ficca et al., 2000; Hoedlmoser et al., 2015). The relationships found between electrophysiologic sleep phenomena (sleep spindles, slow oscillations, REM theta rhythm) and memory are also inconsistent (Casey et al., 2016; Genzel et al., 2009; Landsness et al., 2009; Weigenand et al., 2016; Fogel et al., 2007; Prehn-Kristensen et al., 2013). So, the question of whether sleep’s role in memory consolidation is passive (through reduced interference) or active (through sleep-specific neural processes) remains open. Thus, to gain a clearer understanding of the extent to which sleep itself contributes to improved performance and what is the impact of different sleep stages in this improvement, further investigation is needed.

In our study, we aimed to investigate whether the reduction of either SWS or REM sleep during one night affects the performance on procedural and declarative memory tasks and whether changes in performance correlate with changes in sleep architecture and electrophysiologic sleep phenomena (sleep spindles, slow oscillations, and REM theta rhythm). For the reduction of target sleep stages we used a method of selective suppression by presenting an auditory tone of increasing intensity. Unlike studies in which for SWS and REM disruption participants were woken from these sleep stages (Casey et al., 2016; Genzel et al., 2009; Morgenthaler et al., 2014), we avoided waking the participants entirely whenever possible to make the procedure less stressful. We measured changes in sympathovagal balance and salivary cortisol to control possible stress reactions to sleep disruption.

## 2. Materials and Methods

### 2.1. Participants

A total of 28 healthy volunteers, males, aged from 19 to 27 (23.20 ± 0.45 years (mean ± SEM)) took part in the study. The participants were medication free, abstained from caffeine, chocolate and alcohol 24 h prior to participation. All of them were native Russian speakers. All of them were undergraduate students of Lomonosov Moscow State University. As it follows from their sleep diaries and actigraph data, they slept about 8 hours (7.73 ± 0.18 min; range 7-9 h) per night 1 week before each experiment. The study was performed according to the Declaration of Helsinki on research involving human participants. The ethics committee of the Institute of Higher Nervous Activity and Neurophysiology of the Russian Academy of Sciences approved the research protocol (№ 5, 21 November 2016).

Before the first experiment has started, a written informed consent was obtained, and the participants were allowed to withdraw at any time. All of them were paid for their participation: after each experiment they were paid 1000 RUB (13.67 USD).

### 2.2. Procedure

Each participant took part in at least two experimental sessions: a session with selective SWS suppression during one night’s sleep and a session with regular night sleep as a control condition. Fifteen out of 28 volunteers participated in another session with disturbed sleep: a session with selective REM suppression. The sessions were arranged in a counterbalanced order in both the 28-person and 15-person samples and were separated by 1-4 weeks.

Subjects were unaware whether their sleep would be suppressed or not, but prior to each night (including the control one) they were cautioned that throughout the night they might hear sounds and do not need to react to them.

Participants arrived at the laboratory at 7-8 p.m. and left at 9-10 a.m. the next day. They went to bed in a sleep chamber between 11 p.m. and midnight and were awakened after exactly 8 hours. At night, about 01:30 AM and 04:00 AM, we woke them to take saliva samples for cortisol analysis, such awakenings did not last longer than 5 minutes. After participants have been waken up in the morning, all of them took a glucose tolerance test which was a part of another research. Three hours prior to scheduled night sleep, the lights in the laboratory were dimmed to 10

Lux as bright light could disrupt falling asleep. A sleep chamber was completely dark and soundproof. Except for the sleep interference, the design of all sessions was identical and the participants received the same instructions. The approximate experimental schedule is shown in Table 1.

**Table 1.**
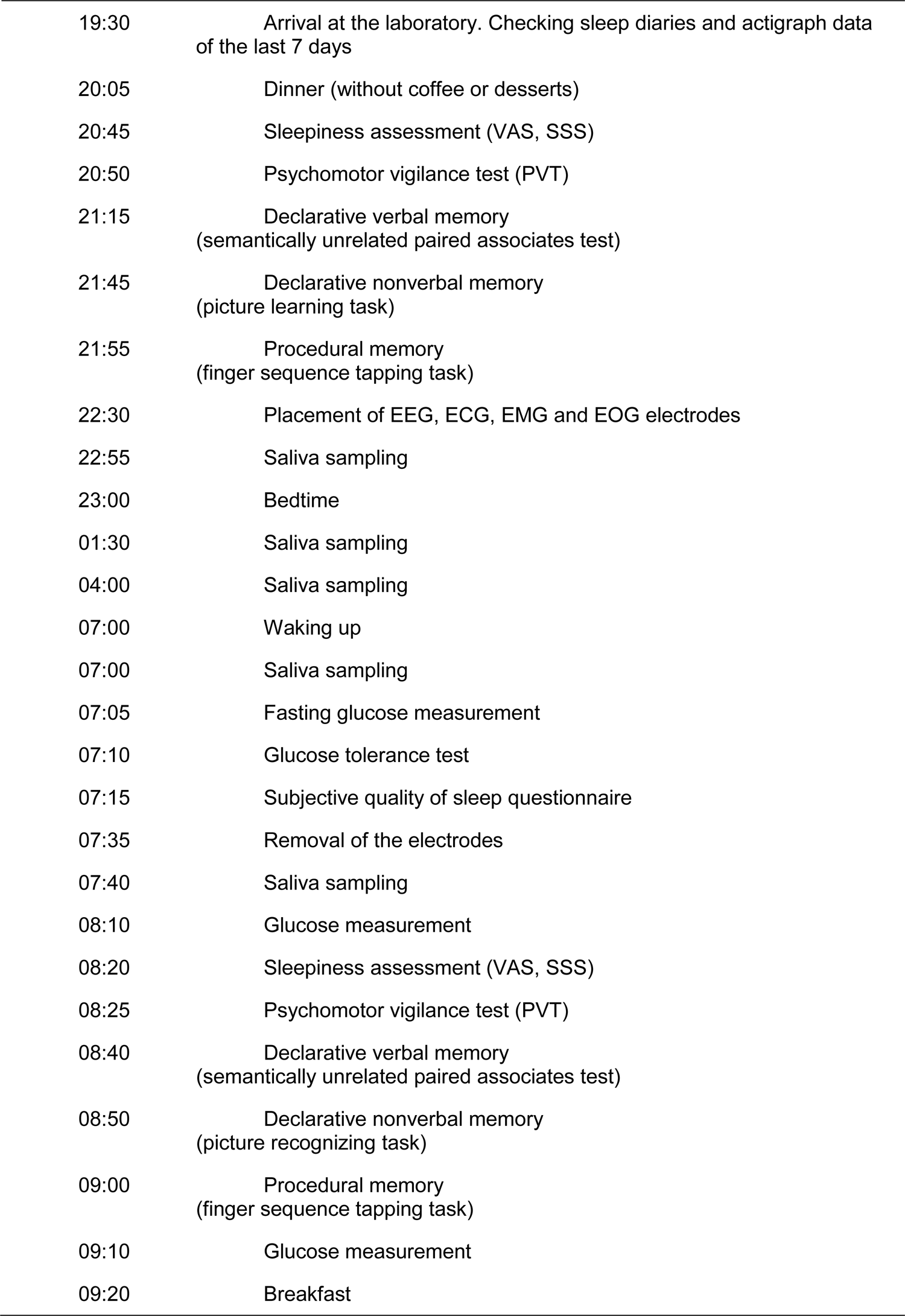
The experimental schedule.

|  |  |
| --- | --- |
| 19:30 | Arrival at the laboratory. Checking sleep diaries and actigraph data of the last 7 days |
| 20:05 | Dinner (without coffee or desserts) |
| 20:45 | Sleepiness assessment (VAS, SSS) |
| 20:50 | Psychomotor vigilance test (PVT) |
| 21:15 | Declarative verbal memory<br>(semantically unrelated paired associates test) |
| 21:45 | Declarative nonverbal memory<br>(picture learning task) |
| 21:55 | Procedural memory<br>(finger sequence tapping task) |
| 22:30 | Placement of EEG, ECG, EMG and EOG electrodes |
| 22:55 | Saliva sampling |
| 23:00 | Bedtime |
| 01:30 | Saliva sampling |
| 04:00 | Saliva sampling |
| 07:00 | Waking up |
| 07:00 | Saliva sampling |
| 07:05 | Fasting glucose measurement |
| 07:10 | Glucose tolerance test |
| 07:15 | Subjective quality of sleep questionnaire |
| 07:35 | Removal of the electrodes |
| 07:40 | Saliva sampling |
| 08:10 | Glucose measurement |
| 08:20 | Sleepiness assessment (VAS, SSS) |
| 08:25 | Psychomotor vigilance test (PVT) |
| 08:40 | Declarative verbal memory<br>(semantically unrelated paired associates test) |
| 08:50 | Declarative nonverbal memory<br>(picture recognizing task) |
| 09:00 | Procedural memory<br>(finger sequence tapping task) |
| 09:10 | Glucose measurement |
| 09:20 | Breakfast |

### 2.3. Materials

#### The subjective sleep assessment questionnaire

After waking up, all the participants completed the questionnaire on subjective sleep quality. The latter consisted of three scales and required to evaluate the depth and quality of sleep from 0 to 10 and estimate the number of nighttime awakenings.

#### Sleepiness assessment instruments

In the evening and in the morning, right before the cognitive tasks we measured the sleepiness of our participants with Stanford Sleepiness Scale (SSS) <u>(Hoddes et al., 1973)</u> and Visual Analogue Scale (VAS). The SSS is a single-item 7-point self-report questionnaire that measures an individual’s subjective level of sleepiness. The VASS consists of two statements with opposite meanings (sleepy and alert) located at the ends of a 100-mm line. The participants were asked to put a vertical mark on the line between these statements at a point that best reflected their perceived alertness.

#### The psychomotor vigilance task

After the assessment of sleepiness, subjects proceeded with the psychomotor vigilance task (PVT) (Dinges, Powell, 1985) included in the PEBL 2.0 software. PVT measures sustained attention, or vigilance, and is widely used in sleep deprivation experiments (Drummond et al., 2005). A fixation cross appeared on a black screen for 400 ms, followed by a target stimulus (red circle, 300 ms) at randomized intervals. The task required hitting the “space” button on a keyboard as soon as a target stimulus appeared. The time interval between two subsequent stimuli altered from 2 to 12 seconds, and the whole task lasted 10 minutes (from 80 to 96 target stimulus presentations). The percentages of correct responses, lapses (i.e., response latency exceeding 500 ms) and false starts (i.e., responses faster than 150 ms) as well *as averaged response time* were calculated. Vigilance was tested equally before and after sleep, whereas declarative and procedural memory was formed in the evening and was tested in the morning.

#### Learning tasks

After PVT the memory encoding (in the evening) or testing (in the morning) followed. Declarative verbal memory was encoded and tested using a **Semantically Unrelated**

**Paired Associates Test** (Tucker, Fishbein, 2008). 120 Russian words all of them nouns with 3 syllables) were combined in 60 pairs so that no semantic relationships appeared. In the evening each pair was presented for 5 sec with an interstimulus interval of 100 msec. To improve the encoding the participants were instructed to visualize two words interacting with each other. For example, to learn a pair of words “lily - desert”, one could imagine a lily growing among the sand. Learning was immediately followed by a cued recall test of 30 pairs from 60, in which the left word in pair (cue) was presented and the right word (target) had to be recalled. Subjects had unlimited time for recalling of the target word, after the response was entered the correct answer was displayed for 2 s. Thus, a half of word pairs was presented for encoding twice. During the morning cued recall test the participants recalled all 60 pairs and no feedback was provided. An order of pairs during each testing was random.

For analysis, the total number of correct responses during morning recall was calculated and also the percentage (with reference to the total number of correct responses) of correct responses separately for the word pairs that were learned once and those that were learned twice. To assess the baseline encoding capacity at the beginning of each experiment, the number of correctly recalled pairs during cued recall of 30 pairs in the evening was calculated.

For testing of Declarative nonverbal memory we used a **Picture Learning Task** (Antonenko et al., 2013). In the evening 50 pictures of landscapes or houses were presented to the participants. They were instructed to memorize them and indicate whether the landscape was tropical or not, or the house was residential or not. For both of options they had to press a left or right key on a keyboard. In the morning of the retrieval 100 pictures were presented, 50 of them were new and 50 pictures had been previously seen. Subjects had to answer if they had already seen the picture by pressing one of four buttons: yes, maybe, maybe not, and no.

For analyses, the first two and the last two types of response, respectively, were pooled. Then the following proportions (with reference to the total number of responses) of pictures correctly recognized by participants were calculated: a proportion of correctly remembered pictures and a proportion of correctly rejected pictures.

To assess procedural memory the **Finger Sequence Tapping Task** (Walker et al., 2002) was used. In this test the participants learned to type on a keyboard a five-digit sequence (e.g. 4–2–3–1–4) with their non-dominant hand. They were instructed to do it as accurately and quickly as possible while using their four fingers (excluding the thumb). After waking up they performed the same task. We expected to see improvement in speed and number of correctly typed sequences. During the learning, subjects were asked to perform 12 30-s blocks with 30-s breaks in between. During the retrieval, they were asked to perform three 30-s blocks. The sequence was presented continuously on a screen. No immediate feedback was given on pressing a key, but after each block the number of correct sequences and the total number of tapped sequences were presented.

For analysis, we used the results of the last three learning blocks performed in the evening and all morning retrieval blocks. At first, the number of correctly tapped sequences averaged across three blocks was calculated. Then the accuracy of tapping was calculated as a proportion of correctly tapped sequences referenced to the total number of tapped sequences. There were three sets of stimuli for memory tasks (five-digit sequences, sets of pictures and word pairs) that were alternated in three sessions and balanced in order across subjects.

#### Polysomnography recording

The EEG was recorded according to the international 10-20 electrode system (Jasper, 1958) using 19-channel recorder Encephalan-EEGR-19/26 and “Encephalan” software (Medicom MTD, Taganrog, Russia) from F3, Fz, F4, C3, Cz, C4, P3, Pz, P4, O1, O2. referenced against mastoid electrodes A1 and A2. We also recorded chin electromyogram (EMG), left and right electrooculogram (EOG), and electrocardiogram in standard lead-II (ECG). The sampling frequency was 250 Hz.

While the subject was sleeping, we monitored the polysomnogram (PSG) in real time. This was done during the control night to ensure the correct software functioning and during experimental nights to selectively suppress SWS or REM sleep. We suppressed the given sleep stage with acoustic tones of increasing intensity until markers of lighter stages (NREM1 or NREM2) appeared in EEG. Waking the participant completely was avoided whenever possible. The tones were played by two speakers located about 30 cm away from the participant’s head. Sound intensified in increments of 5 dB, starting with 35 dB and maximum up to 90 dB. The frequency of sound was 532 Hz. For controlling the tones, a script written in E-Prime was used (E_Prime 1.2, Psychology Software Tools, Pittsburgh, PA, USA).

We assessed sleep stages by standard criteria (Rechtschaffen, Kales, 1968) and guidelines of American Academy of Sleep Medicine (Carskadon, Dement, 2011). SWS start was identified when the delta-rhythm index was higher than 20% (more than 6 seconds of >75 mV and <2 Hz waves in a 30-second epoch). REM sleep was identified when we observed desynchronization in EEG with low EMG amplitude and absence of sleep spindles or K-complexes. Rapid eye movements in EOG and irregular ECG rhythm were used as additional markers of REM, however, we took into account that in tonic REM sleep, rapid eye movements are absent. Primary EEG electrodes that we used for the monitoring were С3, Сz and С4.

**Saliva sampling** and LC-MS/MS **cortisol analysis** were described elsewhere (Ukraintseva et all., 2020).

### 2.4. Data analysis

#### Polysomnography

Two experts identified sleep stages independently using standard criteria (Rechtschaffen, Kales, 1968; Carskadon, Dement, 2011) blinded to test performance results. Inter-scorer reliability was >93%.

We calculated the following sleep parameters: sleep onset latency; wakefulness after sleep onset, WASO; sleep period time, SPT (time after sleep onset before the final awakening); total sleep time, TST (sleep period time without night wakefulness periods); percentage (from TST) of NREM1, NREM2, SWS and REM. Sleep efficiency was calculated as a relation of TST to SPT. For each night we built hypnograms and calculated the number of full sleep cycles. Also, we calculated the number of disturbing sounds presented during suppression nights.

#### EEG analysis

We analyzed EEG of 15 subjects participated on all three sessions: control, with SWS and REM suppression using Brainvision Analyzer software (Brain Products GmbH, Munich, Germany).

All recordings were filtered with a notch-filter (50 Hz), high frequency >0.5 and low-frequency <30 Hz filters. Then we manually removed exogenous and endogenous artifacts, if they were scattered and didn’t cover a significant portion of the recording. In the latter case we performed independent component analysis (ICA) and deleted artifact components via inverse ICA.

We segmented recordings based on sleep stages markers and then broke them into equal segments of 8 seconds. To avoid zero padding we changed sampling rate to 256 Hz, so that the number of data points in each segment became a power of two (8*256=2048). We then performed EEG power spectral analysis in consecutive 8-second epochs (fast Fourier transform routine, Hanning window) for C3, Cz and C4 channels in ten frequency bands: delta1 (0.5–2 Hz), delta2 (2–4 Hz), theta1 (4–6 Hz), theta2 (6–8 Hz), alpha1 (8–10 Hz), alpha2 (10–12 Hz), sigma1 (12– 14 Hz), sigma2 (14–16 Hz), beta1 (16–20 Hz) and beta2 (20-30 Hz). In each recording we averaged power spectrums for four sleep stages: NREM1, NREM2, SWS and REM. Then we averaged spectral power in C3, Cz and C4 and analyzed it statistically.

Detection of sleep spindles was performed in two stages. At the first stage, each electrode was processed separately. We performed continuous wavelet transformation using the Morlet mother wavelet and Matlab 2021a software (Mathworks [https://www.mathworks.com/help/matlab/index.html?s_tid=CRUX_topnav]). The signal power in the spectral range of 10-16 Hz was summarized by frequency to assess spectral power dynamics in time. Spectral threshold of 225 mkV^2 was considered as the beginning of a sleep spindle. This number was obtained empirically for the whole dataset by two experts-somnologists. Sleep spindles separated by a gap of less than 250 ms were merged into one. Sleep spindles lasting less than 300 ms were then removed.

In the next step, sleep spindles that overlapped in time for several electrodes were grouped into one event. The duration of a multi-electrode spindle was considered to be the time interval including all spindles in the group. The spindle frequency was defined for the electrode with the maximal wavelet spectral power.

For each segment of NREM2 and SWS separately we analyzed the number of spindles, their duration, amplitude, frequency and density (the number of spindles referenced to the total duration of NREM2 or SWS per night).

Heart rate variability analysis was described elsewhere (Ukraintseva et all., 2020). Heart rate, LF and HF were calculated. The LF/HF ratio was used to estimate sympathetic modulation.

#### Statistical analysis

Statistical analysis was performed using Statistica 10 software for Windows 7 (Stat Soft. Inc., Tulsa, USA). To assess the distribution type of data collected we used Kolmogorov–Smirnov test. Then parametric T-test and ANOVA (with Newman-Keuls test for post-hoc) for normally distributed data and non-parametric Friedman ANOVA (with Wilcoxon test for post-hoc) for asymmetrically distributed data were applied to the data obtained. Spearman correlation analysis was conducted to assess the correlations between PVT data, procedural and declarative performance and sleep variables. Statistical significance was set at p < 0.05.

## 3. Results

### Sleepiness, polysomnographic data and subjective sleep assessment

Sleep suppression resulted in a reduction of mean duration of SWS from 101,93 to 51,16 min, i.e., by 50% (from 101,63 to 47,60 min, i.e., by 53% if analyzing data of all 28 subjects participated in control and SWS suppression sessions) and REM duration from 89,5 to 43,60 min, i.e., by 51% in respecting suppression experiments. Percentages (from TST) of those stages in the opposite experiments (SWS during REM suppression and vice versa) did not change significantly (Table 2). As expected, the percentage of the NREM1 significantly increased during both suppression nights (p = 0,001 for SWS suppression and p < 0,001 for REM suppression). In the case of slow-wave sleep suppression, in addition, the percentage of NREM2 was higher than in control night (p < 0,001). Though we tried not to awaken the subjects during sleep suppression, the percentage of WASO increased significantly in REM suppression night (p = 0,003). On SWS suppression night, we did not observe significant changes in WASO; only when analyzing data from the entire sample of 28 subjects an upward trend (p = 0,080) was found.

**Table 2.** Mean values of polysomnographic data and subjective assessments of sleep and sleepiness in control session and in sessions with SWS and REM suppression.

|  | N=15 |  |  |  |  | N=28 |  |  |  |
| --- | --- | --- | --- | --- | --- | --- | --- | --- | --- |
|  | Control | P<br>Control /<br>SWS suppression | SWS suppression | P<br>SWS suppression /<br>REM suppression | REM suppression | P<br>Control /<br>REM suppression | Control | P | SWS suppression |
| **Total sleep time (min)** | 412,60± 29,84 | 0,261 | 403,63±41,43 | **0,007** | 365,07±60,36 | **0,001** | 421,88±29,90 | 0,116 | 412,18±38,60 |
| **Sleep period time (min)** | 450,47±18,65 | 0,516 | 444,73±31,08 | 0,983 | 444,93±24,18 | 0,521 | 457,63±18,12 | 0,445 | 453,66±26,24 |
| **WASO (%)** | 9,42±6,17 | 0,275 | 10,54±7,18 | **0,006** | 25,15±22,77 | **0,003** | 8,76±5,94 | 0,080 | 10,56±7,15 |
| **Sleep efficiency** | 0,92±0,05 | 0,281 | 0,91±0,06 | **0,021** | 27,31±39,48 | **0,021** | 0,92±0,05 | 0,084 | 0,91±0,06 |
| **NREM1 (%)** | 5,35±3,70 | **0,001** | 8,76±4,98 | **0,006** | 14,41±7,12 | **<0,001** | 4,76±3,21 | **<0,001** | 7,98±4,62 |
| **NREM2 (%)** | 47,63±5,69 | **<0,001** | 57,08±6,48 | **<0,001** | 47,88±8,32 | 0,885 | 48,45±5,70 | **<0,001** | 57,13±5,21 |
| **SWS (%)** | 24,55±6,79 | **<0,001** | 11,83±5,43 | **<0,001** | 22,53±6,96 | 0,135 | 24,22±6,65 | **<0,001** | 12,13±4,98 |
| **REM (%)** | 21,23±4,46 | 0,511 | 20,19±6,32 | **<0,001** | 11,83±3,20 | **<0,001** | 21,08±4,55 | 0,602 | 20,62±5,62 |
| **N of cycles** | 3,60±0,74 | 0,824 | 3,67±0,98 | 0,161 | 4,00±0,85 | 0,154 | 3,62±0,74 | 0,691 | 3,71±0,90 |
| **N of sounds** | - | - | 995,14±852,81 | 0,064 | 471,21±538,94 | - | - | - | 997,88±879,72 |
| **Depth_Sub** | 7,98±1,20 | **0,012** | 6,83±1,69 | 0,060 | 5,87±2,29 | **0,004** | 7,71±1,61 | 0,069 | 7,07±1,63 |
| **NWake_Sub** | 2,43±1,07 | **0,012** | 4,13±2,22 | **0,035** | 6,67±3,83 | **0,001** | 2,30±1,19 | **0,026** | 3,43±2,13 |
| **Quality_Sub** | 8,00±1,07 | **0,038** | 7,13±1,85 | **0,011** | 5,93±2,05 | **0,001** | 7,96±1,29 | **0,013** | 7,00±1,92 |
| **VASev** | 4,67±2,38 | 0,259 | 3,86±2,15 | 0,514 | 3,53±1,95 | 0,308 | 4,44±2,16 | 0,753 | 4,15±2,00 |
| **VASmo** | 4,11±2,30 | 0,460 | 3,84±2,03 | 0,410 | 4,09±2,02 | 0,496 | 3,80±2,11 | 0,311 | 3,53±1,92 |
| **SSSev** | 3,28±1,03 | 0,397 | 2,87±1,13 | 0,241 | 3,20±1,21 | 0,972 | 3,04±0,90 | 0,753 | 2,93±1,04 |
| **SSSmo** | 3,21±1,37 | 0,678 | 3,07±1,10 | 0,155 | 3,67±1,23 | 0,308 | 2,93±1,24 | 0,754 | 2,85±1,06 |

Note. Data are mean values ± standard deviation. WASO, wakefulness after sleep onset; SWS, slow-wave sleep; REM, rapid eye movement sleep; N of cycles, number of complete sleep cycles, recorded during the night; N of sounds, number of disturbing sounds, presented during the night; Depth\_Sub, subjective depth of sleep; NWake\_Sub, subjective number of nighttime awakenings; Quality\_Sub, subjective sleep quality; VASev, VASmo, Visual Analogue Scale evening and morning values; SSSev, SSSmo, Stanford Sleepiness Scale evening and morning values. Significant differences are highlighted in bold.

The number of disturbing sounds presented during sessions with sleep suppressions did not differ significantly, although, during REM suppression it tended to be lower than during SWS suppression (p = 0,064). We found no between-session difference in the number of complete sleep cycles recorded throughout the night.

According to Friedman ANOVA subjective assessment of the quality of sleep, its depth and the number of awakenings showed significant session effect (Chi Sqr. (N = 15, df = 2) = 16,82 p<0,001 for quality, Chi Sqr. (N = 15, df = 2) = 10,72 p = 0,005 for depth and Chi Sqr. (N = 15, df = 2) = 16,44 p<0,001 for the number of awakenings). The sleep quality and depth were assessed to be lower after two nights with disturbed sleep than after the control one, and the subjective number of awakenings was rated to be higher, respectively (Table 2). At the same time, in the session with REM suppression the sleep quality was the lowest and the number of awakenings was the highest; they differed significantly not only from the control session but also from the session with SWS suppression.

Nevertheless, according to Stanford visual scale and Visual analogue scale results morning and evening sleepiness levels were similar in all sessions.

### The psychomotor vigilance task data

The evening PVT data in three sessions did not differ significantly, suggesting that at the beginning of memory encoding the vigilance level of the participants was similar in all conditions.

The percentage of lapses in PVT increased in the morning after both the control (p = 0.002, N = 28) and REM suppression (p = 0.020, N = 15) nights, data not shown.

Respectively, the percentage of correct responses decreased (p = 0.009, N = 28 after control night and p = 0.011, N =15 after REM suppression night, Fig. 1A). The same goes for the reaction time in these two sessions: it was significantly slower in the morning (p = 0.004, N = 28 after control night and p = 0.045, N = 15 after REM suppression night, Fig 1B).

**Figure 1.**
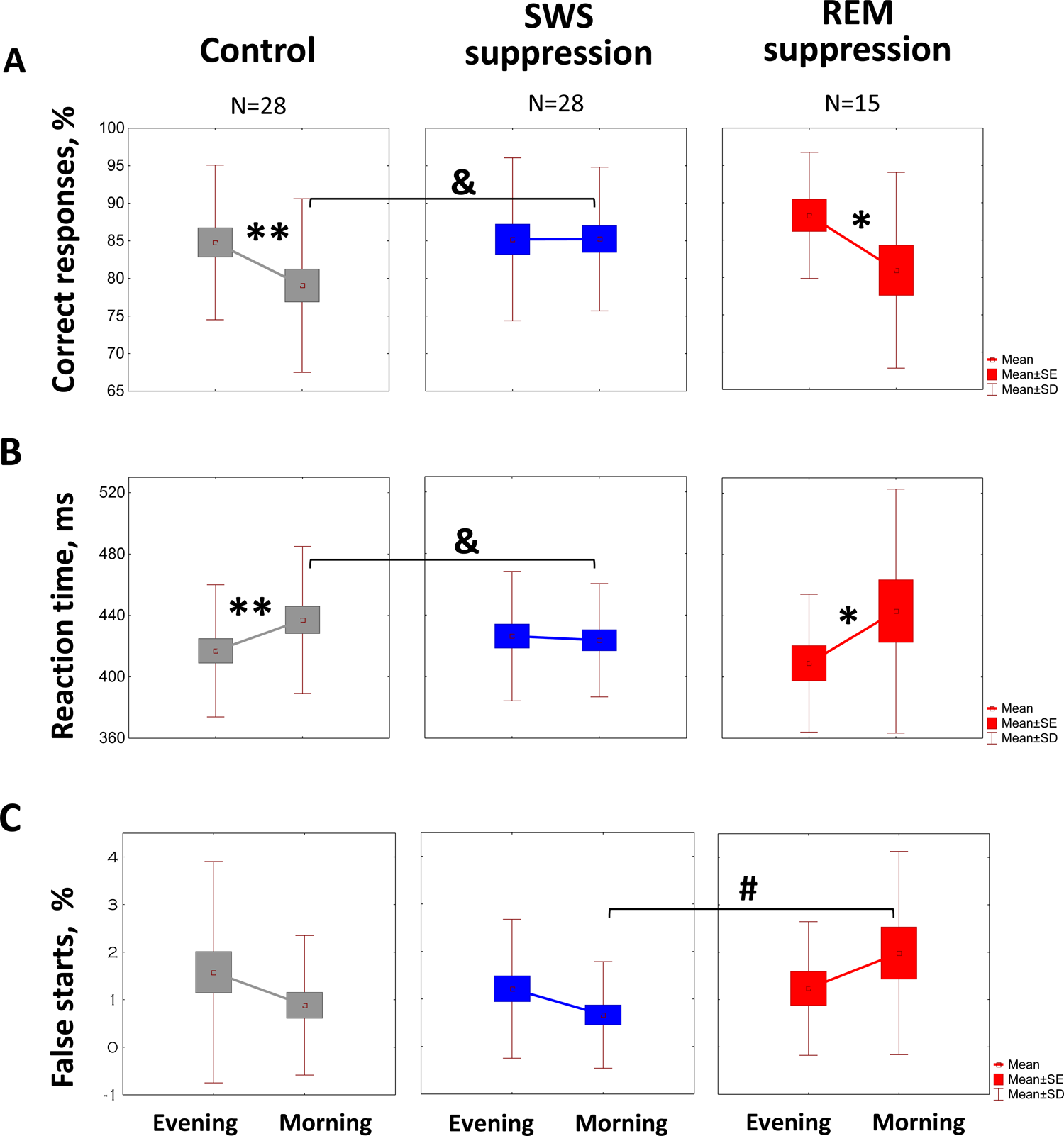
The Psychomotor Vigilance Task performance in the evening and in the morning in control (grey boxes), SWS suppression (blue boxes) and REM suppression (red box) conditions. (A) The percentage of correct responses. (B) The reaction time. (C) The percentage of false starts. *, **, indicates a significant within session difference, p < 0.05, p < 0.01, respectively. &, indicates a significant difference between control and SWS suppression conditions, p < 0.05. #, indicates a significant difference between SWS and REM suppression conditions, p < 0.05.

Surprisingly, such an overnight slowdown in reactions was not detected after the SWS suppression night: the percentages of lapses and correct responses, as well as the reaction time, remained at the same level in the morning. Moreover, in comparison with the control session after SWS suppression there are more correct responses (p = 0.001, N = 28), fewer lapses (p = 0.004, N = 28) and reaction time was faster (p = 0.046, N = 28) indicating higher vigilance level. Comparing with REM suppression session there are fewer false starts (p = 0.035, N = 15, Fig 1C). The comparison of morning PVT data in control and REM suppression sessions (N = 15) did not show significant differences.

### Memory consolidation

#### Verbal declarative memory: Word pairs

The number of correct responses during the evening cued recall of 30 word pairs did not differ between sessions indicating similar baseline encoding ability. The total number of correct responses during morning recall of all 60 word pairs did not differ as well, suggesting that neither SWS nor REM suppression affected memory consolidation. However, when we analyzed the percentage of correct responses separately for word pairs that were learned once and those learned twice, we found a significant session effect: F(2, 28) = 5,2215, p = 0,012. A post hoc analysis showed a significant difference (p = 0,009, N = 15) between two sessions with sleep suppression. The recall rate for word pairs learned twice was higher after the night with SWS suppression (Fig 2A left panel), whereas after the night with REM suppression the participants better recalled pairs learned only once (Fig 2B left panel). A comparison of control and SWS suppression sessions did not reveal any difference in the number or percentage of correctly recalled pairs even when the data of all 28 subjects were analyzed (Fig 2 right panels).

**Figure 2.**
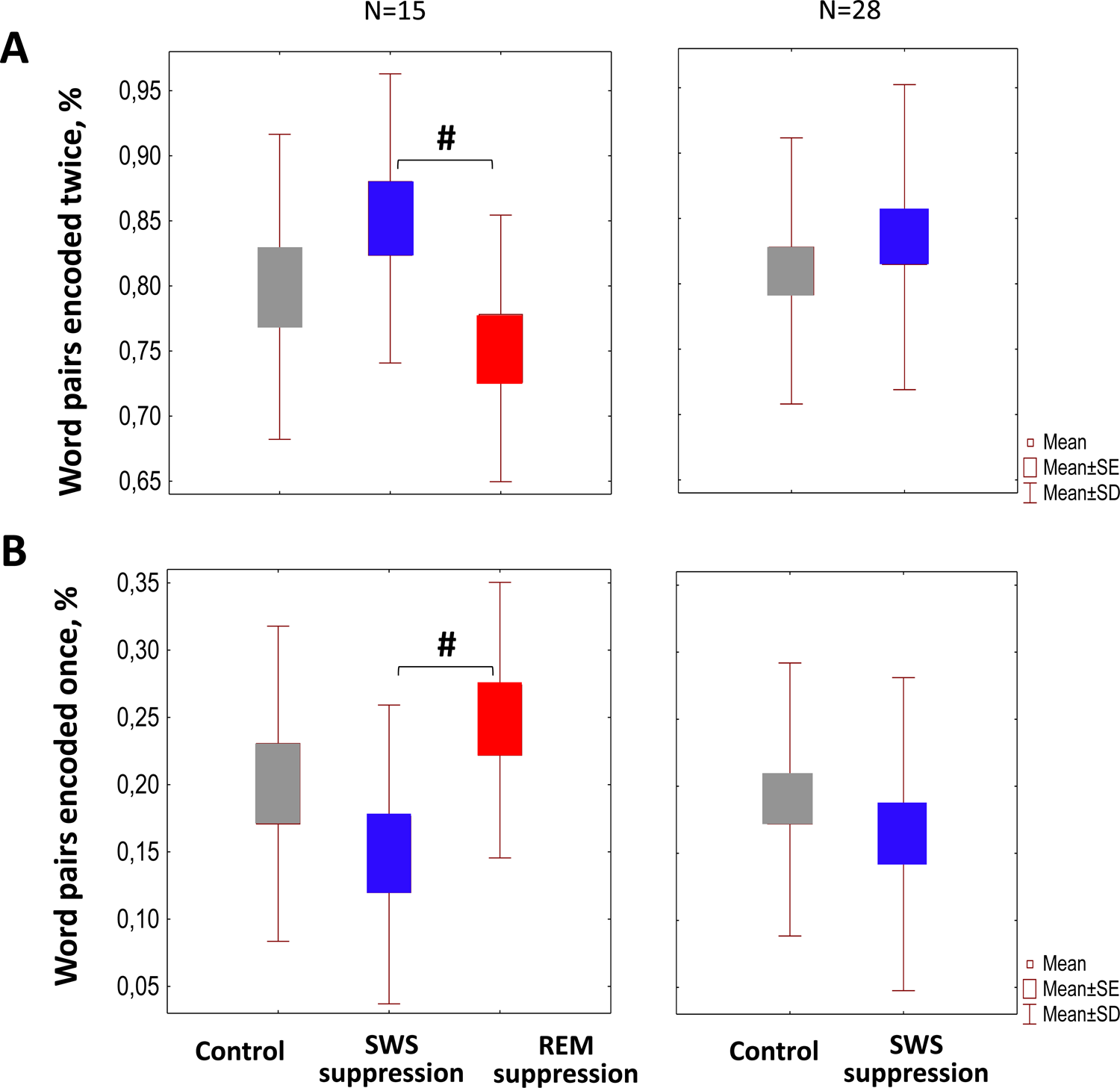
The percentages of word pairs correctly remembered in the morning in control (grey boxes), SWS suppression (blue boxes) and REM suppression (red box) conditions. (A) Word pairs encoded twice during the learning. (B) Word pairs encoded once during the learning. #, indicates a significant difference between SWS and REM suppression conditions, p < 0.05. On the left panels the data of the entire sample (N = 28) in control (grey boxes) and SWS suppression (blue boxes) condition are shown.

#### Non-verbal declarative memory: Picture learning task

The number of pictures correctly recognized by participants in the morning was similar after all three nights spent in the laboratory. No significant differences were found between sessions with disturbed and intact sleep either in the proportion of correctly remembered pictures or in the proportion of correctly rejected pictures.

#### Procedural memory: Finger sequence tapping task

Performance on the finger sequence tapping task, as indicated by the number of correctly tapped sequences per 30-s interval, did not differ between three sessions, either during the evening training or at the morning retrieval testing (Fig. 3A). In all conditions significant overnight improvement was shown (p<0,001 for all conditions; N=28 for control night and night with SWS suppression; N=15 for night with REM suppression). Likewise, the accuracy of tapping grew overnight in all sessions including both with sleep suppression (p<0,001 for all conditions; N=28 for control night and night with SWS suppression; N=15 for night with REM suppression, Fig. 3B).

**Figure 3.**
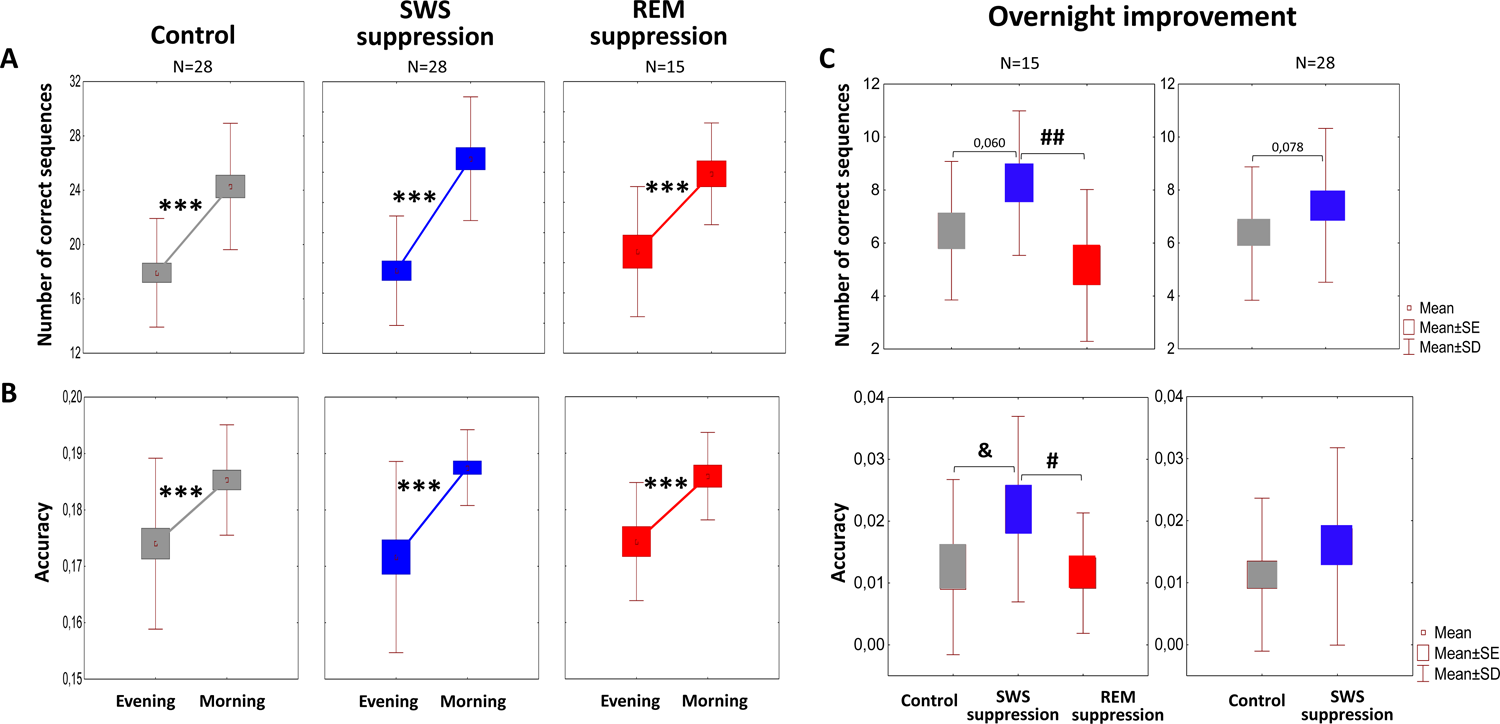
The Finger Tapping Task performance in control (grey boxes), SWS suppression (blue boxes) and REM suppression (red box) conditions. (A) The number of correctly tapped sequences. (B) The accuracy of tapping. Left panels, the performance in the evening and in the morning. Right panels, the overnight improvement. ***, indicates a significant within session difference, p < 0.001. ##, indicates a significant difference between SWS and REM suppression conditions, p < 0.01. The other designations are as in Figure 1 and 2.

However, when we analyzed the overnight improvement of performance (the difference between evening and morning values) with ANOVA for repeated measures, we found statistically significant session effect: F (2, 28) = 5,81, p = 0,008 for the number of correct sequences; and F(2, 28) = 3,77, p = 0,035 for the accuracy. Surprisingly, the participants showed the highest improvement after the night with SWS suppression. A post hoc Newman-Keuls test indicated that the increase in the number of correct sequences was significantly greater than in the session with REM suppression (p = 0.006, N = 15) and tended to be greater than for the control one (p = 0,060, N = 15, Fig. 3C top left panel).

Analyzing data of all 28 subjects participated in control and SWS-suppression sessions, we have also found similar trend for the number of correct sequences (p = 0,078, N = 28, Fig. 3C top right panel).

Likewise, after the night with SWS suppression, the highest overnight increase in accuracy was found (Newman-Keuls test: p = 0,048, N = 15 comparing with REM-suppression night; and p = 0,032, N = 15 comparing with control one, Fig. 3C bottom left panel). However, the comparison of accuracy improvement on a sample of 28 participants did not show significant differences between control and SWS-suppression sessions (Fig. 3C bottom right panel).

### Cortisol data

According to Friedman ANOVA, morning cortisol data showed a significant session effect: Chi Sqr. (N = 15, df = 2) = 8,13 p = 0,017 for 7:00 and Chi Sqr. (N = 15, df = 2) = 6,53 p = 0,038 for 7:40. A post-hoc analysis revealed a decreased salivary cortisol concentration at 07:00 h, i.e., immediately after awakening, in both SWS and REM suppression sessions. There was a significant difference compared with the control for SWS suppression (p = 0,001) and marginally significant (p=0,061) for REM suppression, Fig 4A left panel). After the night with SWS suppression, cortisol remained lowered even 40 minutes after waking up (p = 0,020, in comparison both with control and REM suppression sessions), whereas after REM suppression at this time, it has already returned to control values. A decrease of salivary cortisol after night with SWS suppression was likewise confirmed by an analysis of the entire sample: both at 7:00 and 7:40, it was lower than in the control condition (p = 0,026 for 7:00 and p = 0,011 for 7:40, N = 28, Fig 4A right panel).

**Figure 4.**
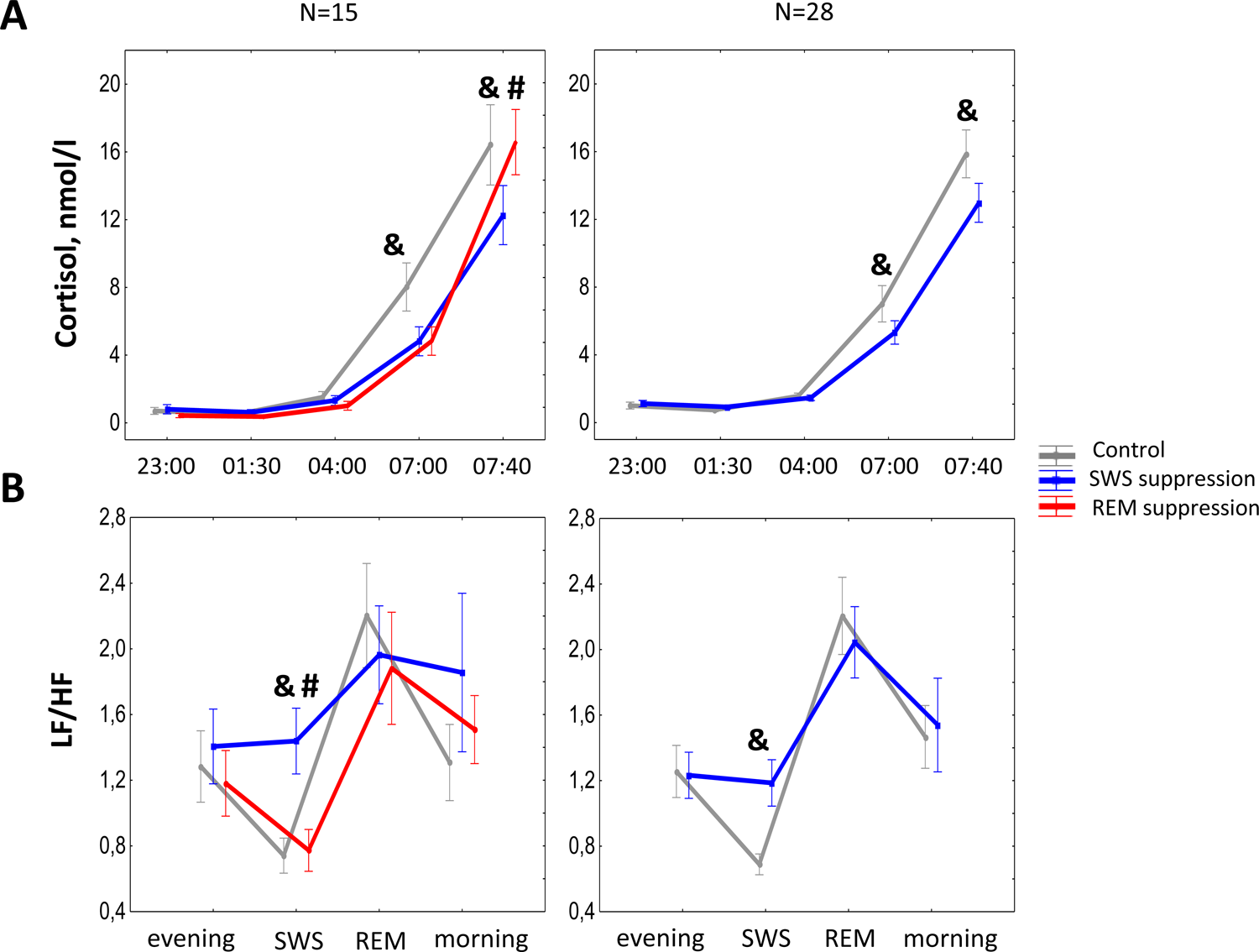
Salivary cortisol (A) and heart rate variability parameter LF/HF(B) in control (grey line), SWS suppression (blue line) and REM suppression (red line) conditions. The other designations are as in Figure 1.

### Heart rate variability data

Friedman ANOVA found a significant session effect on HRV index LF/HF during SWS: Chi Sqr. (N = 15, df = 2) = 10,13 p = 0,006. A post-hoc analysis showed (Fig. 4B left panel) that SWS suppression led to a rise in LF/HF (p = 0,006 in comparison with control and p = 0,003 in comparison with REM suppression sessions). Increased LF/HF was found only during SWS suppression and was not observed neither during subsequent REM sleep nor in the morning after waking up. Likewise, within the entire sample we also found an increase in LF/HF during SWS suppression compared with undisturbed SWS in control condition (p = 0,001, N = 28, Fig. 4B right panel). There were no significant effects of SWS or REM suppression on the other analyzed parameters: heart rate, LF, and HF.

### EEG spectral power

#### SWS suppression effects

According to the results of the Friedman ANOVA test (Chi Sqr. (N = 15, df = 2) = 6,93 p = 0,031) during SWS periods remained in the recordings despite suppression, delta1 spectral power was significantly lower than during undisturbed SWS in control (p = 0,036) and REM suppression (p = 0,031) sessions (Fig. 5 A). At the same time the powers in theta2, alpha1, alpha2, sigma1, sigma2, beta1, and beta2 frequency bands were significantly higher. For theta2 Chi Sqr. (N = 15, df = 2) = 6,93 p = 0,031; p = 0,002 in comparison with control and p = 0,011 in comparison with REM suppression condition. For alpha1 Chi Sqr. (N = 15, df = 2) = 12,40 p = 0,002; p = 0,002 in comparison with control and p = 0,006 in comparison with REM suppression session. For alpha2 Chi Sqr. (N = 15, df = 2) = 10,00 p = 0,007; p = 0,006 in comparison with control and p = 0,003 in comparison with REM suppression session. For sigma1 Chi Sqr. (N = 15, df = 2) = 12,40 p = 0,002; p = 0,001 in comparison with control and p = 0,008 in comparison with REM suppression session. For sigma2 Chi Sqr. (N = 15, df = 2) = 10,53 p = 0,005; p = 0,001 in comparison with control and p = 0,002 in comparison with REM suppression session. For beta1 Chi Sqr. (N = 15, df = 2) = 8,13 p = 0,017; p = 0,002 in comparison with control and p = 0,001 in comparison with REM suppression session. For beta2 Chi Sqr. (N = 15, df = 2) = 8,40 p = 0,015; p = 0,011 in comparison with control and p = 0,015 in comparison with REM suppression session.

**Figure 5.**
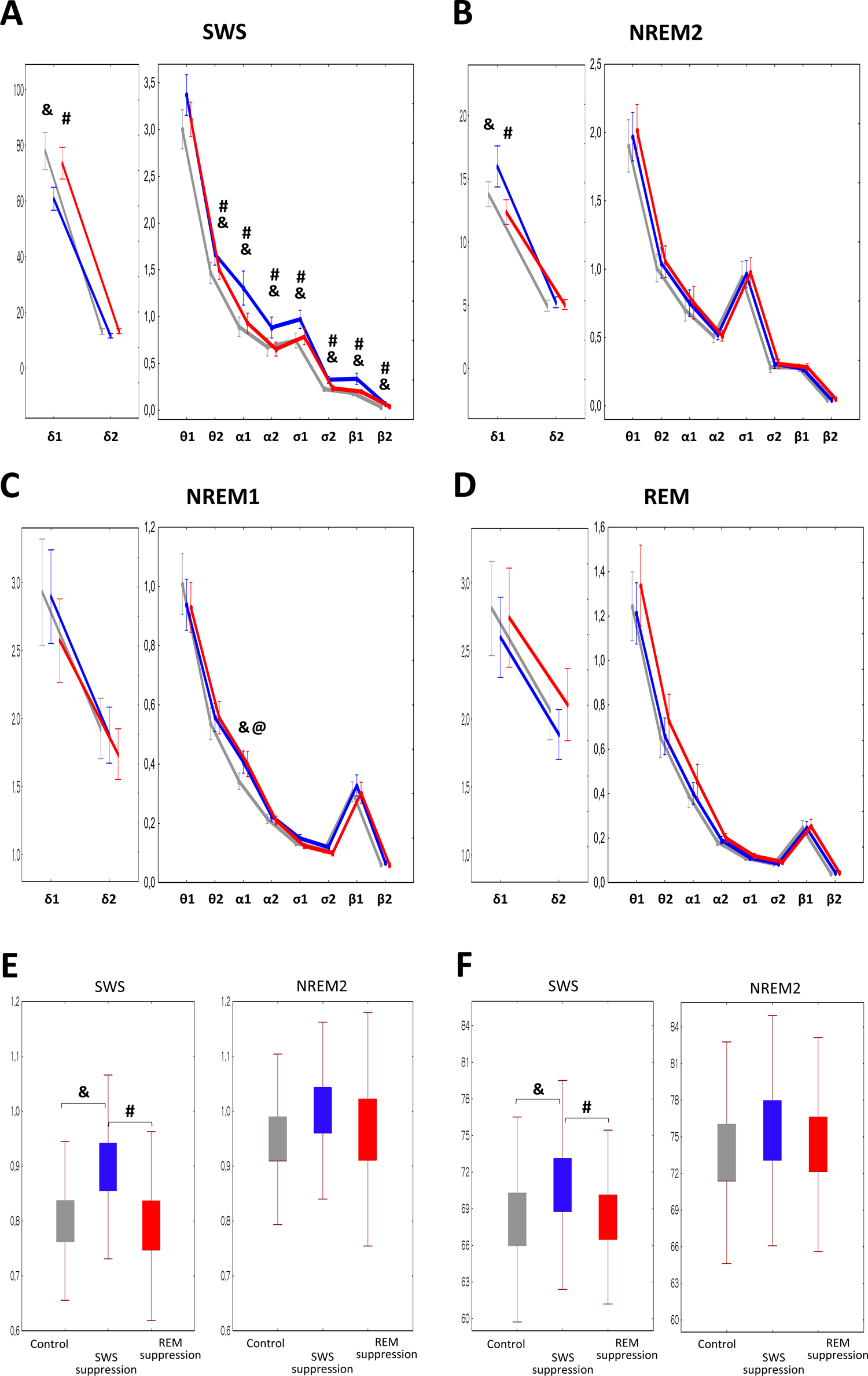
EEG power spectra during SWS (A), NREM2 (B), NREM1 (C) and REM (D) as well as SWS and NREM2 spindle features (duration (E) and amplitude (F)) in control (grey line), SWS suppression (blue line) and REM suppression (red line) conditions.@, indicates a significant difference between control and REM suppression conditions, p < 0.05. The other designations are as in Figure 1.

SWS suppression also affected NREM2 and NREM1 (Fig.5 B, C). During NREM2, unlike SWS, it caused an increase in delta1 activity (Chi Sqr. (N = 15, df = 2) = 10,80 p = 0,005; p = 0,027 in comparison with control and p = 0,015 in comparison with REM suppression condition. During NREM1 an increase in alpha1 spectral power was also found (Chi Sqr. (N = 15, df = 2) = 10,00 p = 0,007; p = 0,008 in comparison with control session).

#### REM suppression effects

EEG power spectra in REM episodes found in PSG after REM suppression did not differ significantly from those for undisturbed REM (Fig. 5 D). However, REM suppression the same as SWS suppression affected NREM1 spectra, causing an increase in alpha1 power (Fig. 5 C; p = 0,011 compared with control night).

### Spindle measures

Analysis of spindle features showed no effect of SWS suppression on NREM2 spindle parameters (Fig. 5 EF right panels). At the same time spindles in SWS periods remained after suppression differed significantly from those in undisturbed SWS (Fig. 5 left panels). They were longer (Chi Sqr. (N = 15, df = 2) = 12,40 p = 0,002; p < 0,001 compared with control and p = 0,005 compared with REM suppression condition) and had higher amplitude (Chi Sqr. (N = 15, df = 2) = 8,40 p = 0,015; p = 0,005 compared with control and p = 0,017 compared with REM suppression condition). The density and frequency of SWS spindles did not differ significantly between sessions.

### Correlation analysis results

To identify the relationships between vigilance, memory retention, and sleep features (sleep architecture and sleep spindles measures as well as delta1 and delta2 spectral power during NREM sleep and theta1 and theta2 power during REM sleep) Spearman correlation analysis was performed.

In the control condition, a positive correlation was found with WASO for the percentage of correct responses in PVT (R = 0,57, p = 0,026) and negative correlations – for the percentage of lapses and reaction time (R = −0,56, p = 0,028 and R = −0,55, p = 0,034, respectively).

The accuracy of tapping in the morning was positively associated with the amount of NREM2 (R = 0,63, p = 0,013) and negatively – with those of SWS (R = −0,74, p = 0,002).

The overnight improvement in accuracy was associated with the duration of SWS spindles (R = 0,73, p = 0,002) and with the amplitude of NREM2 spindles (R = 0,55, p = 0,034). The overnight improvement in the number of correctly tapped sequences was also positively correlated with the duration of SWS spindles (R = 0,54, p = 0,039).

In SWS suppression condition significant correlations were found only with procedural performance: the accuracy of tapping in the morning was positively associated with TST (R = 0,58, p = 0, 025) and negatively – with WASO (R = 0,54, p = 0, 038). The number of correctly tapped sequences in the morning showed a positive correlation with the amplitude of NREM2 spindles (R = 0,57, p = 0,027).

In REM suppression condition the percentage of correct responses in PVT was negatively associated with TST (R = −0,52, p = 0,046) and positively – with WASO (R = 0,68, p = 0,005). The number of correctly tapped sequences in the morning was positively correlated with sleep efficiency (R = 0,72, p = 0,003).

## 4. Discussion

Our data show that the reduction of mean duration of either SWS or REM by more than 50% does not impair memory consolidation. Verbal and nonverbal declarative memory consolidated successfully in both control and suppression sessions and did not respond with deterioration to sleep interference, even though 53% SWS suppression and 51% REM sleep suppression were achieved.

Procedural memory in the finger tapping task also consolidated successfully in all three conditions, showing beneficial effect of night sleep on speed and accuracy of tapping. Surprisingly, after the night with selective SWS suppression an overnight improvement in the skill was even better pronounced than after the other two nights. There were significant differences with REM suppression session for speed and accuracy. Comparing with the control condition this effect was less consistent, but nevertheless, in the sample of 15 participants the difference was significant for the improvement in accuracy; for the improvement in speed we found marginally significant differences both in the sample of 15 and 28 subjects.

Moreover, such paradoxical beneficial effect of SWS suppression also showed the PVT performance. After the control and REM suppression nights, morning PVT data indicated a decline in vigilance: the reaction time slowed, proportion of correct responses decreased, and number of lapses increased. Only after suppression of SWS we did not observe an overnight slowdown in reactions. In addition, comparing with REM suppression session there were less false starts. Based on the found higher vigilance specifically after SWS suppression, we can speculate that these effects could be explained by decreased sleep inertia. It is well known that sleep can have inertia, which impairs cognitive and sensory-motor performance immediately after awakening, and its intensity depends on the depth and duration of SWS (Trotti, 2017). Since testing began approximately an hour and a half after awakening, the effects of sleep inertia could still affect the performance at least in those experiments where SWS was left intact. Correlations between sleep data and morning

PVT scores show a positive association of high vigilance level with WASO and a negative one with TST, further indicating the vulnerability of alertness to sleep inertia that can be caused by prolonged, undisturbed sleep.

Apparently, by reducing the duration and depth of SWS through its selective suppression, without expecting it we imposed a beneficial effect on some activities. Thus, our data support the notion that the depth and duration of SWS can influence vigilance and suggest that this effect may last more than an hour and affect procedural performance.

Our data also show that 50 percent of SWS or REM sleep that remained after suppression is sufficient for successful performance in procedural and declarative tasks in the morning. This result is in line with data obtained earlier that overnight memory retention was not impaired even after a significant disruption of sleep architecture (Genzel et al., 2009). Regarding the effects of REM and SWS suppression on memory consolidation, we found in the literature the conflicting results, with some studies pointing to a negative effect (Casey et al., 2016; Landsness et al., 2009) while others showing no effect of these sleep stages on memory (Genzel et al., 2009; Morgenthaler et al., 2014).

When we analyzed the correlations between sleep data and memory retention we found that, in general, long and undisturbed sleep benefits the consolidation of procedural skills. But at the same time, we found a positive association between the accuracy of tapping in the morning with the amount of NREM2 and a negative one with the amount of SWS. Several researchers believe that NREM2 is more significant for memory consolidation than SWS and REM. At least studies of simple motor tasks like finger tapping have shown such task performance to be correlated with the amount of stage 2 sleep (Genzel et al., 2009; Walker et al., 2002) and to be adversely affected by stage 2 sleep disruptions (Smith, MacNeill, 1994).

There is consistent evidence for the involvement of NREM2 sleep spindles in motor (Fogel, Smith, 2006; Nishida, Walker, 2007) and declarative (Genzel et al., 2009) learning. It is also known that sensory stimulation during NREM sleep can trigger sleep spindles (Sato et al., 2007; Andrillon et al., 2017), leading to an increase in their density (Landsness et al., 2009) and suggesting a possible stimulatory rather than inhibitory effect on memory consolidation of such sleep interference. Since we used acoustic stimulation for sleep suppression, we also observed an elevated spindle activity during SWS suppression which was manifested in both sigma band spectral power and the duration and amplitude of the spindles in SWS.

We found significant positive correlations between morning performance in procedural task and sleep spindle features. However, associations with SWS spindles were observed only in control condition, whereas in SWS suppression condition we found associations only with NREM2 spindles, which were not affected by SWS suppression.

Thus, our results do not allow us to draw an unambiguous conclusion about what exactly caused the overnight improvement in procedural skill in session with SWS suppression. However, they show that this improvement was not caused by boosting sleep spindle activity through acoustic stimulation.

Another sleep feature considered to be essential for memory consolidation is slow-wave activity (SWA). Reducing SWA during sleep by acoustic stimulation was found to suppress the sleep-dependent improvement in visuomotor tasks (Landsness et al., 2009; Landsness et al., 2011) and also in perceptual learning (Aeschbach et al., 2008). Our data shows that SWS suppression led to a decrease of spectral power in the delta1 (0,5 – 2 Hz) band during SWS periods. At the same time, during NREM2 periods the power of these slow oscillations was significantly increased possibly due to K-complexes evoked by sounds when they coincided with NREM2 periods. However, we did not find any significant correlations between memory retention and SWA power.

Theta oscillations predominated in REM are also demonstrated to be involved in memory consolidation (Fogel et al., 2007; Prehn-Kristensen et al., 2013), however, we did not find any significant correlations between REM theta power and performance in procedural and declarative tasks in the morning.

Regarding non-verbal declarative memory, we failed to find any relationships between sleep architecture or electrophysiologic sleep phenomena and retention of pictures. For verbal declarative memory we observed the difference between sessions with SWS and REM sleep suppression in recall of word pairs learned with and without testing. In general, the words that were only passively reviewed by the subjects before going to bed were recalled significantly worse each morning compared to the words that were actively practiced in the evening. However, after the night with REM suppression the proportion of correct recall of once-learned words was higher than after the night with SWS suppression. Different influence of SWS and REM suppression on consolidation of such weak and strong memory engrams is difficult to interpret, and data on this problem available in the literature are inconsistent (Tucker et al., 2008; Ukraintseva, Dorokhov, 2011; Deliens et al., 2013; McDevitt et al., 2015; Barsky et al., 2015). Some researchers showed beneficial effects of NREM sleep on weak or exposed to interference memory traces (Ukraintseva, Dorokhov, 2011; Deliens et al., 2013), while others point to the crucial role of REM sleep (McDevitt et al., 2015; Barsky et al., 2015).

It should be noted that in this study, investigating the impact of SWS and REM on memory consolidation we tried to avoid such consequences of sleep disturbance as stress and emotional discomfort, as well as endocrine fluctuations associated with them. A standard practice in sleep science is to deprive sleep stages by waking up the participants (Casey et al., 2016; Genzel et al., 2009; Morgenthaler et al., 2014). We used a relatively mild acoustic sleep-perturbation approach and carefully avoided waking the participant whenever possible.

HRV data showed only a transient increase in sympathetic activity directly at the time of SWS suppression; the next morning the sympathovagal balance did not differ significantly from the control values. Cortisol secretion also was not increased in the morning after nights with sleep suppression comparing with the control condition. Thus, we may conclude that we could avoid the influence of stress on the results obtained.

## 5. Limitations

Some limitations of our study should be mentioned. Despite the number of disturbing sounds presented during the both nights with sleep suppressions did not differ significantly and even tended to be lower during REM suppression, REM interference caused a higher increase in WASO which should be taken into account when comparing the effects of these sleep disturbances. Particularly, an increase in WASO may be responsible for the worse subjective assessments of sleep quality obtained after the night with REM suppression. The explanation for this is likely to be found in the divergent characteristics of both sleep stages in terms of EEG-response to acoustic stimuli. There is evidence that the EEG response to acoustic stimulation is the lowest during REM-sleep (Williams et al., 1964). This is also confirmed by the EEG data we obtained, indicating that disturbing sounds during REM suppression did not cause any significant changes in EEG power spectra (Fig. 5D). Therefore, REM is more difficult to disturb than SWS and REM interference is more likely to cause an awakening than a transition to lighter sleep stages. So, REM suppression might be more detrimental to the well-being than SWS suppression.

## 6. Conclusions

In summary, we found that neither vigilance nor memory consolidation were impaired after about 50% SWS or REM sleep suppression. Unexpectedly, a beneficial effect of selective SWS suppression on vigilance was found. Similarly, after a night with SWS suppression, overnight improvement in procedural skill was even higher than after a night with REM suppression and after a night with undisturbed sleep. Thus, provided that the sleep disturbances do not cause stress, half the normal amount of SWS or REM are sufficient for procedural and declarative memory consolidation. Moreover, selective SWS suppression may improve performance, possibly by reducing sleep inertia associated with prolonged, undisturbed sleep. The study findings help to clarify the role of sleep features per se in the development of cognitive deficits in disorders associated with insomnia or other conditions characterized by disrupted sleep.

## Author Contributions

Conceptualization, K.S., Y.U.; methodology, O.T., K.S. and Y.U.; software, O.T.; validation, K.S., Y.U.; formal analysis, K.S., Y.U.; investigation, K.S., Y.U.; data curation, Y.U.; writing—original draft preparation, Y.U.; writing—review and editing, K.S., Y.U. and O.T.; visualization, K.S., Y.U. and O.T.; supervision, Y.U.; project administration, Y.U. All authors have read and agreed to the published version of the manuscript.

## Institutional Review Board Statement

The study was conducted in accordance with the Declaration of Helsinki, and approved by Ethics Committee of the Institute of Higher Nervous Activity and Neurophysiology of the Russian Academy of Sciences (protocol code № 5, 21.11.2016).

## Informed Consent Statement

Written informed consent has been obtained from the subjects to publish this paper.

## Data Availability Statement

The data presented in this study are available on request from the corresponding author.

## Acknowledgments

We gratefully acknowledge the help of the volunteers who participated in this study. We want to thank our colleagues in the human higher nervous activity lab, especially Maria Antonova for technical support.

## References

Aeschbach D, Cutler AJ, Ronda JM. A role for non-rapid-eye-movement sleep homeostasis in perceptual learning // J Neurosci 28: 2766 –2772, 2008.

Andrillon T, Pressnitzer D, Léger D, Kouider S. Formation and suppression of acoustic memories during human sleep // Nat Commun. 2017 Aug 8;8(1):179. doi: 10.1038/s41467-017-00071-z.

Antonenko, D., Diekelmann, S., Olsen, C., Born, J., & Mölle, M. (2013). Napping to renew learning capacity: enhanced encoding after stimulation of sleep slow oscillations. European Journal of Neuroscience, 37(7), 1142–1151.

Backhaus J., Junghanns K. Daytime naps improve procedural motor memory // Sleep Medicine. 2006. V. 7(6). P. 508–512.

Barsky M.M., Tucker M.A., Stickgold R. REM sleep enhancement of probabilistic classification learning is sensitive to subsequent interference // Neurobiol Learn Mem. 2015 Jul:122:63–8. doi: 10.1016/j.nlm.2015.02.015. Epub 2015 Mar 11.

Brodt S., Inostroza M., Niethard N., Born J. Sleep-A brain-state serving systems memory consolidation // Neuron. 2023 Apr 5;111(7):1050–1075. doi: 10.1016/j.neuron.2023.03.005.

Carskadon M.A., Dement W.C. Monitoring and staging human sleep. Principles and practice of sleep medicine // Principles and practice of sleep medicine, 5th edition / edited by M.H. Kryger, T. Roth, W.C. Dement. St. Louis: Elsevier Saunders, 2011. P. 16–26.

Casey S.J., Solomons L.C., Steier J., Kabra N., Burnside A., Pengo M.F., Moxham J., Goldstein L.H., Kopelman M.D. Slow wave and REM sleep deprivation effects on explicit and implicit memory during sleep // Neuropsychology. 2016. V. 30(8). P. 931–945.

Clemens Z., Fabo D., Halasz P. Overnight verbal memory retention correlates with the number of sleep spindles // Neuroscience 2005;132:529–35

Deliens G., Leproult R., Neu D., Peigneux P. Rapid eye movement and non-rapid eye movement sleep contributions in memory consolidation and resistance to retroactive interference for verbal material // Sleep. 2013 Dec 1;36(12):1875–83. doi: 10.5665/sleep.3220.

Dinges D.F., Powell J.W. Microcomputer analyses of performance on a portable, simple visual RT task during sustained operations // Beh Res Meth Instr Comp. 1985. V. 17. P. 652–655.

Drummond, Sean & Bischoff-Grethe, Amanda & Dinges, David & Ayalon, Liat & Mednick, Sara & Meloy, Mj. (2005). The Neural Basis of the Psychomotor Vigilance Task. Sleep. 28. 1059–68. 10.1093/sleep/28.9.1059.

Ficca G., Lombardo P., Rossi L., Salzarulo P. Morning recall of verbal material depends on prior sleep organization // Behav. Brain Res. 2000. V. 112. P. 159 – 163.

Fogel S.M., Smith C.T. Learning-dependent changes in sleep spindles and Stage 2 sleep // J Sleep Res 15: 250 –255, 2006,

Fogel S.M., Smith C.T., Cote K.A. Dissociable learning-dependent changes in REM and non-REM sleep in declarative and procedural memory systems // Behav Brain Res. 2007 Jun 4;180(1):48–61. doi: 10.1016/j.bbr.2007.02.037. Epub 2007 Feb 28.

Genzel L., Dresler M., Wehrle R., Grözinger M., Steiger A. Slow Wave Sleep and REM Sleep Awakenings Do Not Affect Sleep Dependent Memory Consolidation // Sleep. 2009. V. 32(3). P. 302–310.

Hoddes E., Zarcone V., Smythe H., Phillips R., Dement W.C. Quantification of sleepiness: a new approach // Psychophysiology. 1973. V. 10(4). P. 431–436.

Hoedlmoser K., Birklbauer J., Schabus M., Eibenberger P., Rigler S., Mueller E. The impact of diurnal sleep on the consolidation of a complex gross motor adaptation task. // J Sleep Res 2015 24:100 –109.

Izawa S. et al. REM sleep–active MCH neurons are involved in forgetting hippocampus-dependent memories // Science 2019;365(6459):1308–1313.

Jasper H.H. The Ten-Twenty Electrode System of the International Federation // Electroencephalography and Clinical Neurophysiology. 1958. V. 10. P. 371–375.

Lahl O., Wispel C., Willigens B., Pietrowsky R. An ultra short episode of sleep is sufficient to promote declarative memory performance // Journal of Sleep Research. 2008. V. 17(1). P. 3–10.

Landsness E.C., Crupi D., Hulse B.K., Peterson M. J., Huber R., Ansari H., Coen M., Cirelli C., Benca R. M., Ghilardi M. F., Tononi G. Sleep-dependent improvement in visuomotor learning: a causal role for slow waves // Sleep, 2009;32(10):1273–1284.

Landsness EC, Ferrarelli F, Sarasso S, Goldstein MR, Riedner BA, Cirelli C, Perfetti B, Moisello C, Ghilardi MF, Tononi G. Electrophysiological traces of visuomotor learning and their renormalization after sleep // Clin Neurophysiol 2011.

McDevitt E.A., Duggan K.A., Mednick S.C. REM sleep rescues learning from interference // Neurobiol Learn Mem. 2015 Jul:122:51–62. doi: 10.1016/j.nlm.2014.11.015. Epub 2014 Dec 11.

Meule A. Reporting and interpreting working memory performance in n-back tasks // Front. Psychol. 2017. V. 8. P. 352.

Morgenthaler J., Wiesner C.D., Hinze K., Abels L.C., Prehn-Kristensen A., Göder R. Selective REM-Sleep Deprivation Does Not Diminish Emotional Memory Consolidation in Young Healthy Subjects // PLoS ONE. 2014. V. 9(2). P. e89849.

Nishida M, Walker MP. Daytime naps, motor memory consolidation and regionally specific sleep spindles. PLoS One 2: e341, 2007.

Poe G.R. Sleep Is for Forgetting // The Journal of Neuroscience, 2017, 37(3):464 – 473

Prehn-Kristensen A, Munz M, Molzow I, Wilhelm I, Wiesner CD, Baving L. Sleep promotes consolidation of emotional memory in healthy children but not in children with attention-deficit hyperactivity disorder. PLoS One. 2013 May 29;8(5):e65098. doi: 10.1371/journal.pone.0065098. Print 2013.

Rasch B., Born J. About sleep’s role in memory // Physiol. Rev. 2013. V. 93. P. 681–766.

Rechtschaffen A., Kales A. A Manual of Standardized Terminology, Techniques, and Scoring System for Sleep Stages of Human Subjects. Washington DC: US Government Printing Office, 1968. 57 p.

Sara S.J. Sleep to Remember // The Journal of Neuroscience, 2017, 37(3):457– 463

Sato Y., Fukuoka Y., Minamitani H., Honda K. Sensory stimulation triggers spindles during sleep stage 2 // Sleep 2007;30:511–8.

Smith C., MacNeill C. Impaired motor memory for a pursuit rotor task following Stage 2 sleep loss in college students // J Sleep Res 1994;3:206–13.

Trotti L.M. Waking up is the hardest thing I do all day: Sleep inertia and sleep drunkenness // Sleep Med Rev. 2017, 76–84. doi: 10.1016/j.smrv.2016.08.005.

Tucker M.A., Fishbein W. Enhancement of declarative memory performance following a daytime nap is contingent on strength of initial task acquisition // Sleep. 2008 Feb;31(2):197–203. doi: 10.1093/sleep/31.2.197.

Ukraintseva Yu.V., Dorokhov V.B. [Effect of daytime nap on consolidation of declarative memory in humans] [Article in Russian] // Zh Vyssh Nerv Deiat Im I P Pavlova. 2011 Mar-Apr;61(2):161–9

Ukraintseva Yu.V., Liaukovich K.M., Saltykov K.A., et al. Selective slow-wave sleep suppression affects glucose tolerance and melatonin secretion. The role of sleep architecture Sleep Med, 67 (2020), pp. 171–183, 10.1016/j.sleep.2019.11.1254

Walker, M.P., Brakefield, T., Morgan, A., Hobson, J.A. & Stickgold, R. (2002) Practice with sleep makes perfect: sleep-dependent motor skill learning. Neuron, 35, 205–211.

Weigenand A., Mölle M., Werner F., Martinetz T., Marshall L. Timing matters: open-loop stimulation does not improve overnight consolidation of word pairs in humans // Eur J Neurosci. 2016 Oct;44(6):2357–68.

Williams H.L., Hammack J.T., Daly R.L., Dement W.C., Lubin A. Responses to auditory stimulation, sleep loss and the eeg stages of sleep // Electroencephalogr. Clin. Neurophysiol. 1964. 16, 269—279.

